# Phosphatidylinositol 3-phosphate promotes 53BP1 condensate-like assembly at DNA double-strand breaks

**DOI:** 10.64898/2026.04.18.710733

**Authors:** Nan Xiong, Gaofeng Cui, Xuanqi (Emily) Xu, Yawen Xie, Shih-Chieh Ti, Shikang Liang, Viji M. Draviam, Yang Liu, Cheng-han Yu, Georges Mer, Michael Shing Yan Huen

## Abstract

53BP1 nuclear bodies are dynamic structures with properties resembling biomolecular condensates, but the molecular determinants that govern 53BP1 higher-order assembly at DNA double-strand breaks (DSBs) remain to be established. Here, we show that 53BP1 condensation is stimulated by phosphatidylinositol 3-phosphate (PI(3)P) *in vitro* through a highly specific interaction with its C-terminal tandem BRCT domain (tBRCT). Consequently, mutational inactivation of the 53BP1 tBRCT domain compromised PI(3)P binding and suppressed 53BP1 optodroplet formation *in vivo*. We further show that rapid 53BP1 clustering following DNA damage precedes its stable assembly on DSB-flanking chromatin, requires its tBRCT, and is suppressed by sequestration of nuclear PI(3)P. Taken together, our findings identify PI(3)P binding as a mechanism that promotes the formation and maturation of 53BP1 condensate-like assemblies on damaged chromatin.

## Introduction

DNA double-strand breaks (DSBs) are demarcated by the local accrual of DNA damage signaling and repair proteins and can be microscopically visualized as DNA damage foci. The PI3K-like protein kinases, in particular ATM and DNA-PKcs, play an apical role in the formation of DNA damage foci by phosphorylating the histone variant H2AX at Ser139 (γH2AX) in the vicinity of DSBs, setting off the hierarchical assembly of DNA damage response (DDR) proteins, including 53BP1, which in turn underlies effective checkpoint activation and DNA repair processes^1–3^.

53BP1 is a multi-domain ∼220 kDa protein and serves scaffolding roles at DNA damage foci, anchoring key DDR factors that effect non-homologous end joining (NHEJ) and ATM-dependent checkpoint control^4^. While its C-terminal tandem BRCT domain (tBRCT) has phospho-binding activity^5, 6^, 53BP1 tBRCT engages in a number of phospho-dependent as well as phospho-independent interactions, including with p53^7^, MUM1/EXPAND1^8^, USP28^9^, the MRN complex^10^, and γH2AX^11–13^. Intriguingly, via yet-to-be defined mechanisms, the tBRCT domain also appears to affect its multimeric configuration *in vitro*^10^.

53BP1-containing DNA damage foci have emerged as nuclear compartments with properties of biomolecular condensates^14, 15^. 53BP1 harbors an oligomerization domain^16^ and forms higher-order assemblies *in vitro* and *in vivo* in an intrinsically disordered region (IDR)-dependent manner that can be stimulated by RNA^17–19^. This behavior has been linked to p53-dependent gene activation and the choice of DSB repair pathway^19, 20^. 53BP1 phase separation has also been reported to play a role in maintenance of heterochromatin integrity^21^, although it remains unknown how 53BP1 condensation is regulated beyond the requirement for its IDR.

Nuclear inositol lipids have been implicated in cell cycle regulation and early DNA repair processes^22–24^, and more recently documented to enrich at damaged chromatin to drive ATR-dependent DDRs^25^. Inspired by observations that phospholipids can promote biomolecular condensation *in vitro*^26^, in this study we report a highly specific role of phosphatidylinositol 3-phosphate (PI(3)P) in directing the higher-order assembly of the DDR factor 53BP1. Our findings suggest that IDR-dependent 53BP1 condensation is promoted by its tBRCT-PI(3)P interaction.

## Results

### Early 53BP1 nuclear bodies are sensitive to 1,6-hexanediol

53BP1 and γH2AX have been widely used as surrogate markers for DNA double-strand breaks (DSBs). Interestingly, while γH2AX is required for the stable and productive assembly of 53BP1 on DSB-flanking chromatin, early DNA damage foci containing 53BP1 are discernible in H2AX-deficient cells^27^. Consistently, our time-course examination of DNA damage foci in irradiated U2OS cells revealed a sub-population of 53BP1 nuclear bodies that did not overlap with γH2AX signals at early time points (15 min post IR; **Fig. 1a & Extended Data Fig. 1a**). This is in stark contrast to 53BP1 and γH2AX foci which, by 120 min post IR, overlapped extensively (**Fig. 1a**). Arguing against antibody- and cell type-specific effects, similar observations were made in RPE-1 cells that endogenously express knocked-in GFP-fused 53BP1 (RPE-1 ^GFP^53BP1)^28^(**Fig. 1b**).

**Fig. 1.**
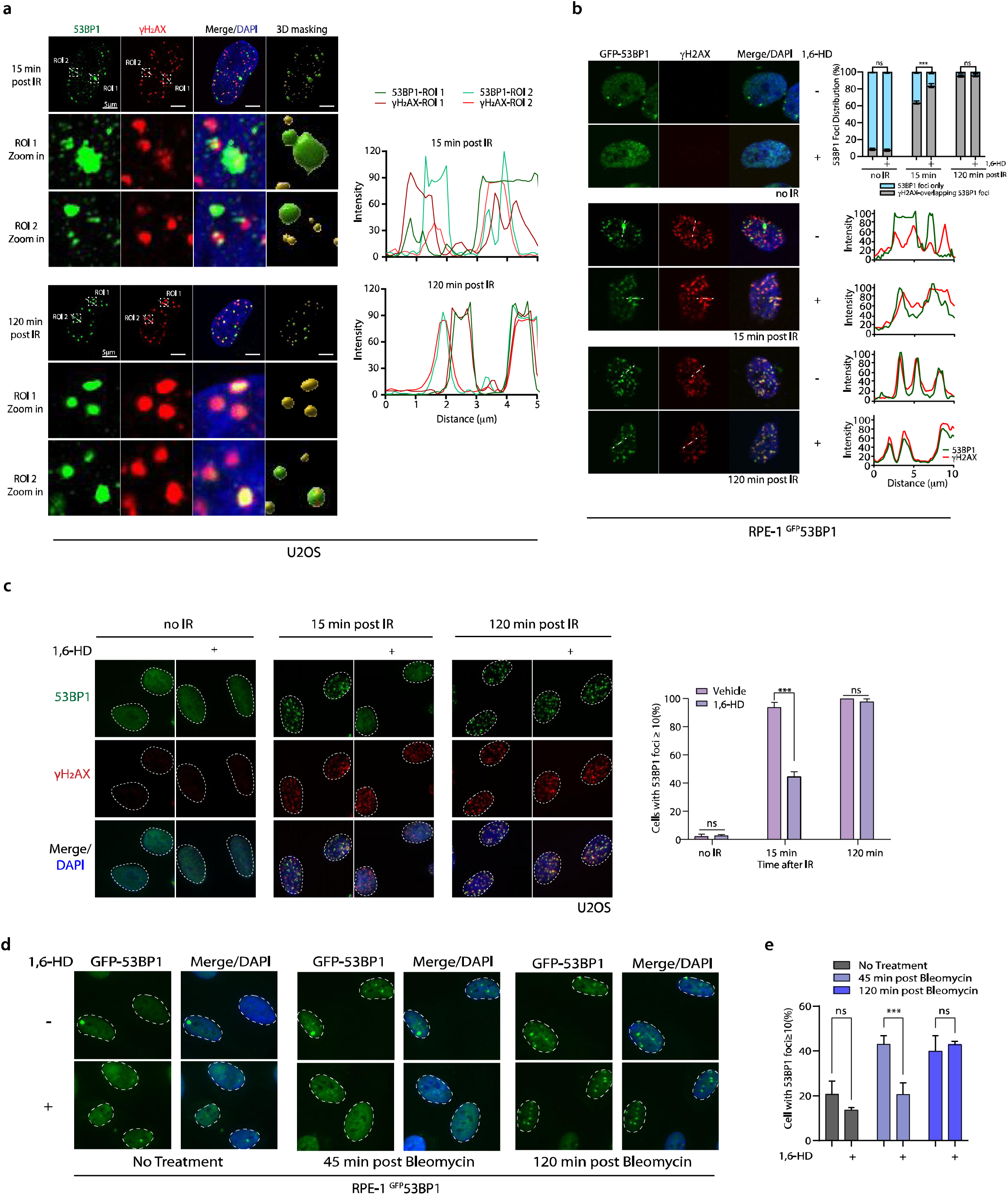
Early 53BP1 nuclear bodies are sensitive to 1,6-HD. **a,** Super-resolution microscopy images reveal maturation of 53BP1 nuclear bodies over time following DNA damage. Cells were fixed and stained with indicated antibodies at 15 and 120 minutes post IR. Fluorescence intensity at specified region of interest (ROI) was quantified and plotted. **b,** Treatment with 1,6-HD eliminated γH2AX-non-overlapping 53BP1 foci in RPE-1^GFP-53BP1^ cells. Cells were irradiated with 3 Gy, permeabilized and pre-treated with 2% 1,6-HD prior to fixation for immunofluorescence at 15 and 120 minutes post IR. The percentages of 53BP1 foci that do or do not overlap with γH2AX were quantified. Fluorescence intensity of 53BP1 and γH2AX signals was measured along the white line and plotted. **c-d,** 1,6-HD treatment dissolved IR-(c) and bleomycin-induced (d) early 53BP1 DNA damage foci. Data represent quantification from three independent experiments (n = 300); bars show mean ± SEM; ***P < 0.001; ns, not significant. Nuclei are outlined with dotted lines.

The idea that 53BP1 forms condensates^17^ prompted us to examine whether early 53BP1 clustering is sensitive to disruption of weak multivalent interactions, including liquid-liquid phase separation (LLPS)^29^. To this end, we incubated triton-permeabilized cells with 1,6-hexanediol (1,6-HD) prior to formaldehyde fixation and examined GFP-53BP1 foci distribution. To minimize potential artefacts, 1,6-HD was applied at 2% (w/v) for only 10 seconds, as described previously^30, 31^. Intriguingly, such 1,6-HD treatment preferentially dissolved γH2AX-non-overlapping 53BP1 nuclear bodies 15 min post treatment with ionizing radiation (IR) (**Fig. 1b**). The time-dependent effect of 1,6-HD on dissolving 53BP1 foci was recapitulated in irradiated U2OS cells (**Fig. 1c**) and in RPE-1 GFP-53BP1 cells challenged with the radiomimetic bleomycin (**Fig. 1d**). Finally, cell pre-treatment with sorbitol, which suppresses 53BP1 foci formation^17^, also preferentially eliminated γH2AX-non-overlapping 53BP1 nuclear bodies (**Extended Data Fig. 1b**).

The above observations led us to hypothesize that 1,6-HD-sensitive 53BP1 nuclear bodies are products of its clustering and condensation in the vicinity of DSBs, and that their formation precedes and is independent of H2AX and its phosphorylation. Indeed, 1,6-HD pre-treatment caused a further marked reduction of 53BP1 foci in H2AX-deficient cells following IR exposure (**Extended Data Fig. 2a**). Together with the discordance between bleomycin-induced GFP-53BP1 foci size and number over time (**Extended Data Fig. 2b**), we propose that early 53BP1 nuclear bodies represent biomolecular condensates, and that their nucleation precedes their stable assembly at DSBs.

### 53BP1 condensation is promoted by PI(3)P *in vitro*

Studies have shown that 53BP1 forms condensates, possibly through LLPS, in a manner that requires its IDR (**Fig. 2a**)^17–19^. Interestingly, during our computational analysis of 53BP1 protein properties^32, 33^, we found that its C-terminal tBRCT domain exhibited a high prediction score for phospholipid binding (**Fig. 2a**). Inspired by the recent description that phospholipids are enriched in biomolecular condensates^26^, we purified a C-terminal foci-forming construct of 53BP1 (residues 1230-1972) (**Fig. 2b**), and observed fusion events characteristic of biomolecular condensation *in vitro* (**Fig. 2c**). To examine whether phospholipids may indeed modulate 53BP1 clustering and higher-order assembly *in vitro*, we incubated phosphatidylinositol (PI) and each of the seven phosphoinositides (PIPs) with purified 53BP1 proteins and assessed 53BP1 droplet formation. Strikingly, we found a highly specific role of PI(3)P in stimulating 53BP1 droplet formation (**Fig. 2d-e**). Supporting a role of tBRCT as a phospholipid-binding domain, deletion (ΔtBRCT) or point mutations that inactivate its phospho-binding activity (R1811A/K1814M) hampered 53BP1 droplet formation *in vitro* (**Extended Data Fig. 3a-c**). Notably, the R1811A/K1814M mutant exhibited a stronger loss-of-function phenotype than ΔtBRCT, as the deletion mutant retained residual droplet-forming capacity. Further, co-addition of RNA had an additive effect on 53BP1 droplet formation (**Extended Data Fig. 3a-c**).

**Fig. 2.**
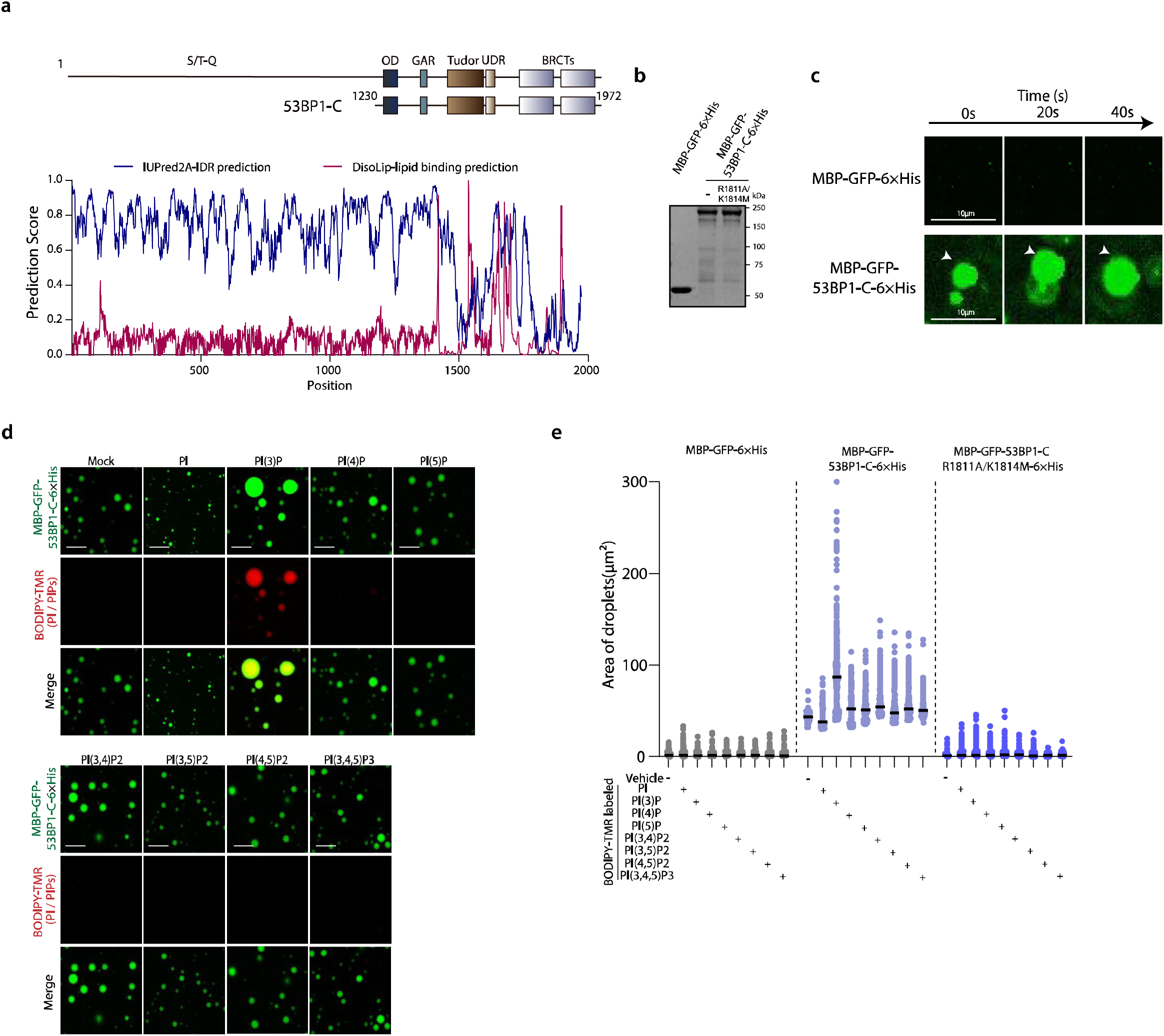
The 53BP1 C-terminal foci-forming region undergoes condensation *in vitro*. **a,** Schematic illustration of conserved 53BP1 domains and prediction scores for intrinsic disorder and lipid binding propensity. **b,** Coomassie blue staining of purified MBP-GFP-6×His, MBP-GFP-53BP1-C-6×His, and MBP-GFP-53BP1-C R1811A/K1814M-6×His proteins. **c,** *In vitro* phase separation assays show that MBP-GFP-53BP1-C-6×His, but not MBP-GFP-6×His, forms fusion droplets in phase separation buffer. Scale bar: 10 µm. **d,** Addition of PI(3)P enhances phase separation of MBP-GFP-53BP1-C-6×His *in vitro*. Mixture contained 5 µM protein and 2.5 µM BODIPY-TMR labeled PI/PIPs. Scale bar: 10 µm. **e,** Quantification of droplet formation across three independent experiments, presented with mean values (solid line).

### 53BP1 tBRCT is a *bona fide* PI(3)P-interacting domain

Given the stimulatory role of PI(3)P in driving 53BP1 condensate formation *in vitro* in a tBRCT-dependent manner, we next examined whether the 53BP1 tBRCT domain can directly interact with PI(3)P. To this end, we prepared giant unilamellar vesicles (GUVs) containing either BODIPY-TMR-labeled PI or PI(3)P, as previously described^34^ (**Fig. 3a**), and incubated them with purified 53BP1 proteins (**Fig. 2b**). Importantly, 53BP1 exhibited robust colocalization with PI(3)P-coated GUVs, but not with PI-coated GUVs (**Fig. 3b**). Moreover, neither of the 53BP1 tBRCT mutants (R1811A or K1814M) accumulated at the GUV boundaries, supporting the idea that the C-terminal tBRCT domain of 53BP1 mediates PI(3)P binding.

**Fig. 3.**
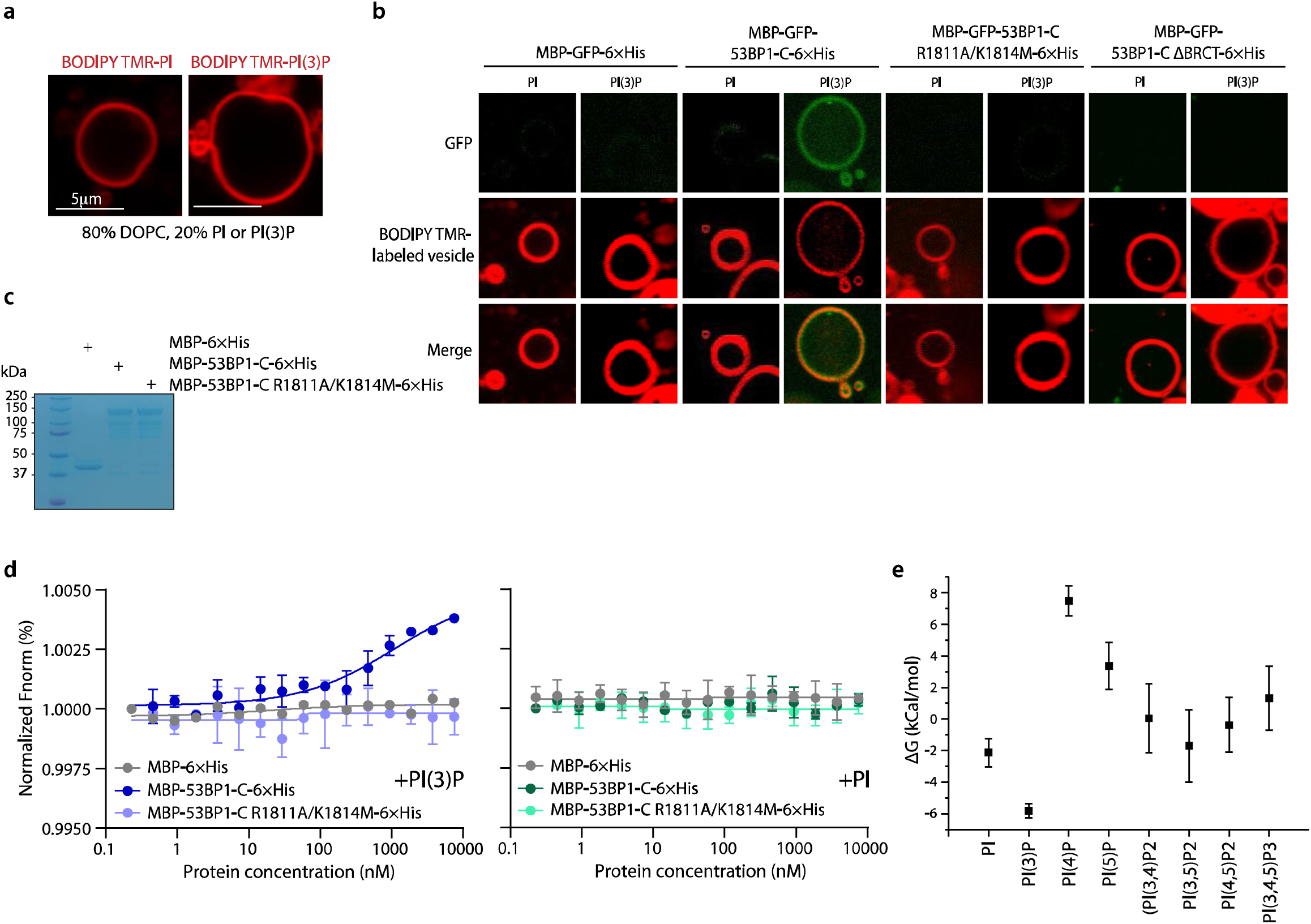
PI(3)P specifically binds to the 53BP1 C-terminal foci-forming region. **a,** GUVs composed of 20% PI or PI(3)P and 80% 1,2-dioleoyl-sn-glycero-3-phosphocholine (DOPC) were prepared. **b,** Accumulation of MBP-GFP-53BP1-C-6×His at PI(3)P-coated GUVs. **c,** Coomassie blue staining of purified MBP-6×His, MBP-53BP1-C-6×His, and MBP-53BP1-C R1811A/K1814M-6×His. **d,** MST binding assays measured the affinity of MBP fusion proteins for labeled PI or PI(3)P, with binding curves plotted to show relative affinity. **e,** Binding free energies of phosphatidylinositol and the indicated phosphoinositides determined by thermodynamic integration using four independent replicates. Error bars represent s.d.

To complement the GUV imaging assay, we further examined 53BP1 tBRCT binding to PI(3)P by microscale thermophoresis (MST) using BODIPY-TMR-labeled ligands. Consistent with a direct interaction, MST experiments revealed reproducible binding between 53BP1 tBRCT and PI(3)P (*K*_d_ ∼ 489 μM, n = 3), but not with PI (**Fig. 3c–d**). In contrast, the R1811A/K1814M mutant showed negligible affinity for PI(3)P. It is important to note that this *K*_d_ reflects a 1:1 binding stoichiometry. In the more relevant context of multivalent interactions between PI(3)P-containing vesicles or clusters and oligomeric 53BP1, the effective binding is much stronger through avidity effects. This is supported by the robust colocalization of 53BP1 C-terminal foci-forming construct with BODIPY-TMR PI(3)P (**Fig. 2d**) and PI(3)P-containing GUVs (**Fig. 3b**).

To determine how 53BP1 tBRCT preferentially interacts with PI(3)P, we carried out atomistic molecular dynamics (MD) simulations and energy calculations with PI and the seven phosphoinositide variants differing in the number and positions of phosphate groups (see Methods; **Extended Data Fig. 4a**). We used AutoDock Vina^35, 36^ to generate plausible docking poses of the different compounds on the tBRCT surface. In the highest-ranked pose for each ligand, the phosphate groups of the variants were positioned within the previously identified phosphate-binding pocket of the 53BP1 tBRCT domain and interact with R1811 and K1814^11^ (**Extended Data Fig. 4b**). Five independent 1 μs MD replicates were then performed using AMBER24^37^ to assess the stability of the best docking poses. Ligand root-mean-square deviations (RMSDs) were calculated after alignment of the 53BP1 tBRCT domain. The RMSD profiles showed that most binding poses were relatively unstable over the course of the simulations (**Extended Data Fig. 4c**). In four of the five trajectories, PI dissociated from the binding pocket, indicating weak binding. In contrast, PI(3)P remained stably bound in all five replicates. The other variants dissociated more readily than PI(3)P during the MD simulations.

To further assess ligand-binding stability and evaluate the docking-derived binding modes, we calculated binding free energies for PI and the seven phosphoinositide species using a stepwise thermodynamic integration (TI) approach^38^ (**Fig. 3e**). Starting structures for TI were taken from well-equilibrated MD simulations. Among all ligands tested, only PI(3)P showed a favourable binding free energy with the 53BP1 tBRCT domain (−5.84 ± 0.44 kcal·mol⁻¹; **Fig. 3e**). These calculations model a single tBRCT domain interacting with a monomeric phosphoinositide ligand. In the relevant multivalent context where oligomeric 53BP1 engages clustered PI(3)P species, multivalent avidity effects are expected to substantially strengthen the interaction. By contrast, the calculated binding free energies for PI(4)P and PI(5)P were positive, indicating unfavourable binding under the modelled conditions. Consistently, both ligands displayed reduced stability in MD simulations, including dissociation in some replicates and large RMSD fluctuations while bound (**Extended Data Fig. 4c**). The calculated free energies for the remaining ligands were close to zero, also consistent with very weak or negligible interaction, in agreement with the large RMSD fluctuations in their MD trajectories.

### Inactivation of 53BP1 tBRCT hampers optodroplet formation *in vivo*

When fused with the blue light-responsive Cryptochrome 2 (Cry2) domain, forced oligomerisation of 53BP1 promotes optodroplet formation *in vivo*^17^. To examine whether 53BP1 tBRCT contributes to biomolecular condensation, we expressed wildtype 53BP1 and its tBRCT mutants in the W1495A background. The W1495A mutation prevents interaction with histone H4 mono- or dimethylated at K20 and releases 53BP1 from chromatin^17, 39^, thereby facilitating optodroplet formation and enabling better analysis of condensation kinetics. Consistent with the proposition that the tBRCT domain promotes 53BP1 condensation, the R1811A/K1814M mutation hampered optodroplet formation when expressed in the context of either full-length 53BP1 or a C-terminal construct that retains foci-formation capability (**Fig. 4a-b**). In line with its ability to form droplets *in vitro*, the ΔtBRCT mutant exhibited only marginal differences compared to the wildtype counterpart in initial formation of optodroplets, similar to a previous report^17^. However, unlike full-length 53BP1, continuous blue-light induction did not promote further clustering of the ΔtBRCT mutant (see 150 seconds), supporting a role for the tBRCT-PI(3)P interaction in driving IDR-dependent optodroplet formation. Our findings highlight a positive role for tBRCT-dependent PI(3)P recognition in promoting 53BP1 condensation and suggest that point mutations in the tBRCT domain may represent a “gain-of-function” phenotype that is distinct from complete domain deletion. Overall, PI(3)P recognition may provide specificity in nucleating the condensation process (see Discussion).

**Fig. 4.**
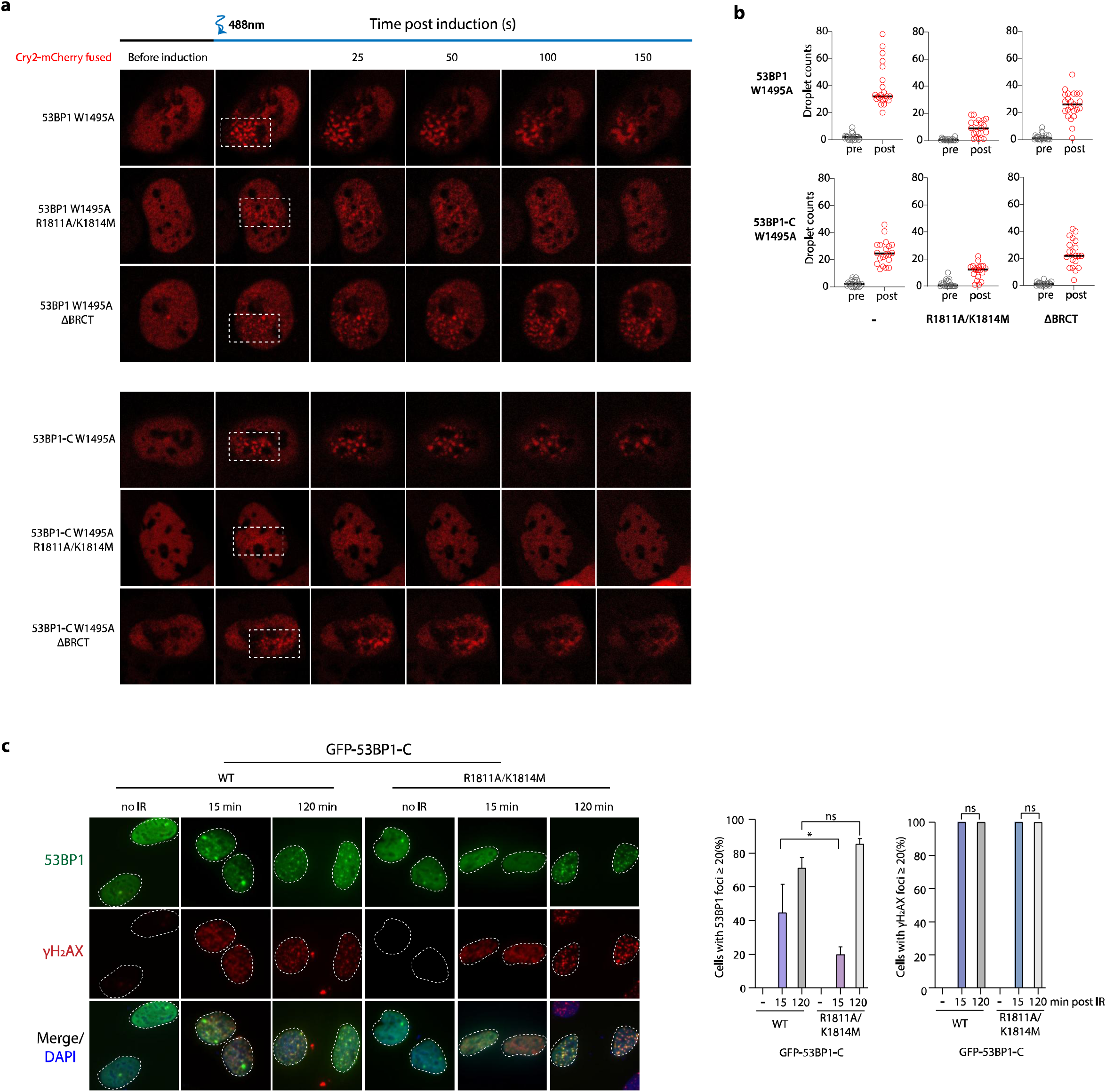
The 53BP1 BRCT R1811A/K1814M mutant impairs assembly. **a,** Cells expressing full-length or C-terminal 53BP1 carrying the R1811A/K1814M mutations exhibited reduced light-induced optodroplet formation compared to wildtype. Representative images before and after light induction are shown. **b,** Quantification from single-cell analysis across three experiments (n≥19); mean ± SEM. **c,** GFP-53BP1-C R1811A/K1814M mutants show fewer DNA damage foci at 15 minutes post IR. U2OS cells transfected with wildtype (WT) or mutant constructs were irradiated, fixed at 15 and 120 minutes, respectively, and stained for γH2AX. Nuclei are outlined with dotted lines. Quantification of three independent experiments (n = 300); bars represent mean ± SEM; *P < 0.05; ns, not significant.

### A 53BP1 tBRCT mutant exhibits delayed kinetics in DNA damage foci formation

That tBRCT binding to PI(3)P 53BP1 promotes condensation prompted us to examine whether this interaction contributes to higher-order assembly during early DSB responses. We reasoned that early 53BP1 DNA damage foci may represent condensate-like assemblies (see **Fig. 1**) that undergo reconfiguration and maturation, facilitating productive accumulation on γH2AX-decorated chromatin domains. To test this idea, we expressed GFP-fused 53BP1 or corresponding R1811A/K1814M mutant and analyzed their ability to form foci following cell exposure to IR. Consistently, point inactivation of the tBRCT domain impaired 53BP1 foci formation at 15 min post-IR (**Fig. 4c**). These observations suggest that the 53BP1 tBRCT domain plays a positive role in promoting condensation during the early stages of the DSB responses.

### Sequestration of nuclear PI(3)P hampers early 53BP1 DNA damage foci formation

Phospholipids have recently been documented to accumulate at DNA damage sites^25^. Consistently, a nuclear localization signal (NLS)-fused GFP-p40PX reporter, which specifically binds PI(3)P, accumulated robustly at laser-induced DNA damage tracks (**Fig. 5a**). Congruent with the established early role of PARylation in the DDR, chemical inhibition of poly(ADP-ribose) polymerase (PARP) activity with olaparib (PARPi) reduced p40PX accumulation at DNA damage sites (**Fig. 5b**). Moreover, cell pre-treatment with the inhibitor of phosphatidyl inositol transfer proteins 1 and 2 (PITP1/2) Osunprotafib (PITP1/2i) also attenuated p40PX recruitment to laser-induced DSBs (**Fig. 5c**). Similarly, inactivation of PI(3)P binding via the R57Q mutation compromised p40PX accrual at laser-induced DNA damage tracks (**Fig. 5d**). Immunostaining further supported PI(3)P enrichment at DSBs (**Fig. 5e**).

**Fig. 5.**
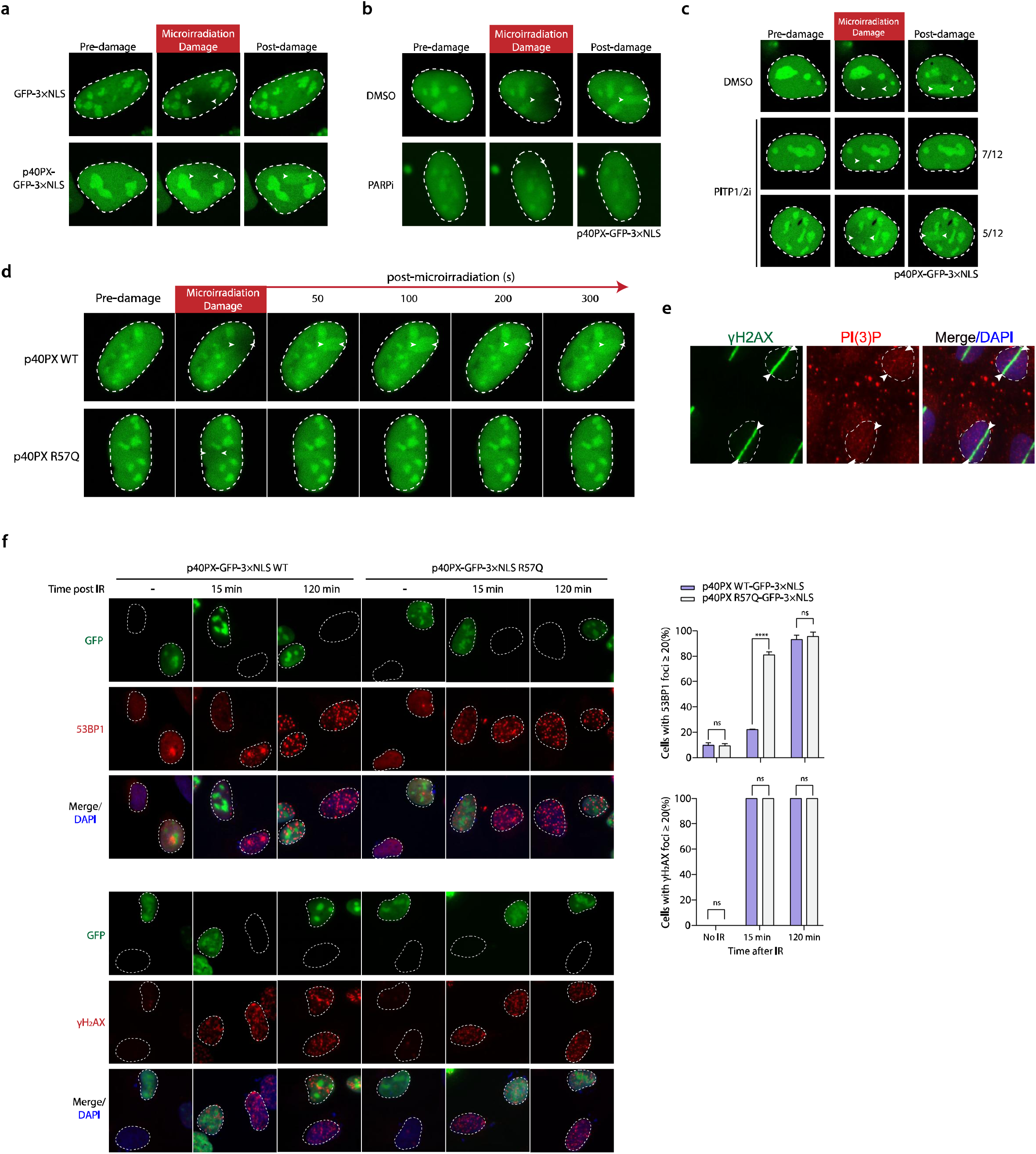
PI(3)P accumulates at DSBs and promotes IR-induced 53BP1 foci formation. **a,** GFP-p40PX-3×NLS (PI(3)P reporter) accumulates at laser-induced DNA damage tracks in U2OS cells, as shown by time-lapse imaging after two-photon laser micro-irradiation. **b,** Inhibition of PARP activity reduces PI(3)P reporter accumulation at DNA breaks. **c,** Inhibition of PITP1 and PITP2 activities hampered PI(3)P reporter accumulation at DNA breaks. **d,** Time-lapse imaging and intensity quantification of cells expressing wildtype (WT) p40PX-GFP-3×NLS or its R57Q mutant following cell exposure to micro-irradiation. **e,** Immunofluorescence with PI(3)P-specific antibody shows co-localization with γH2AX at laser-induced DNA damage sites. **f,** Cells over-expressing p40PX-GFP-3×NLS (WT) or its R57Q mutant were irradiated (3 Gy) and fixed at specified time following IR. Expression of p40PX-GFP-3×NLS correlates with reduced 53BP1 foci formation. Quantification of three independent experiments (n = 300); bars represent mean ± SEM; ****P < 0.0001; ns, not significant.

To further assess whether PI(3)P contributes to early 53BP1 condensation events, we overexpressed p40PX to sequester the nuclear pool of PI(3)P and examined 53BP1 foci formation following IR. In strong support of a positive role of PI(3)P in facilitating swift formation of 53BP1 higher-order assemblies, overexpression of p40PX, but not the PI(3)P-binding-deficient R57Q mutant, impaired foci formation (**Fig. 5f**). Together, these findings suggest that the tBRCT-PI(3)P interaction promotes 53BP1 condensation at DSBs (**Fig. 6**).

**Fig. 6.**
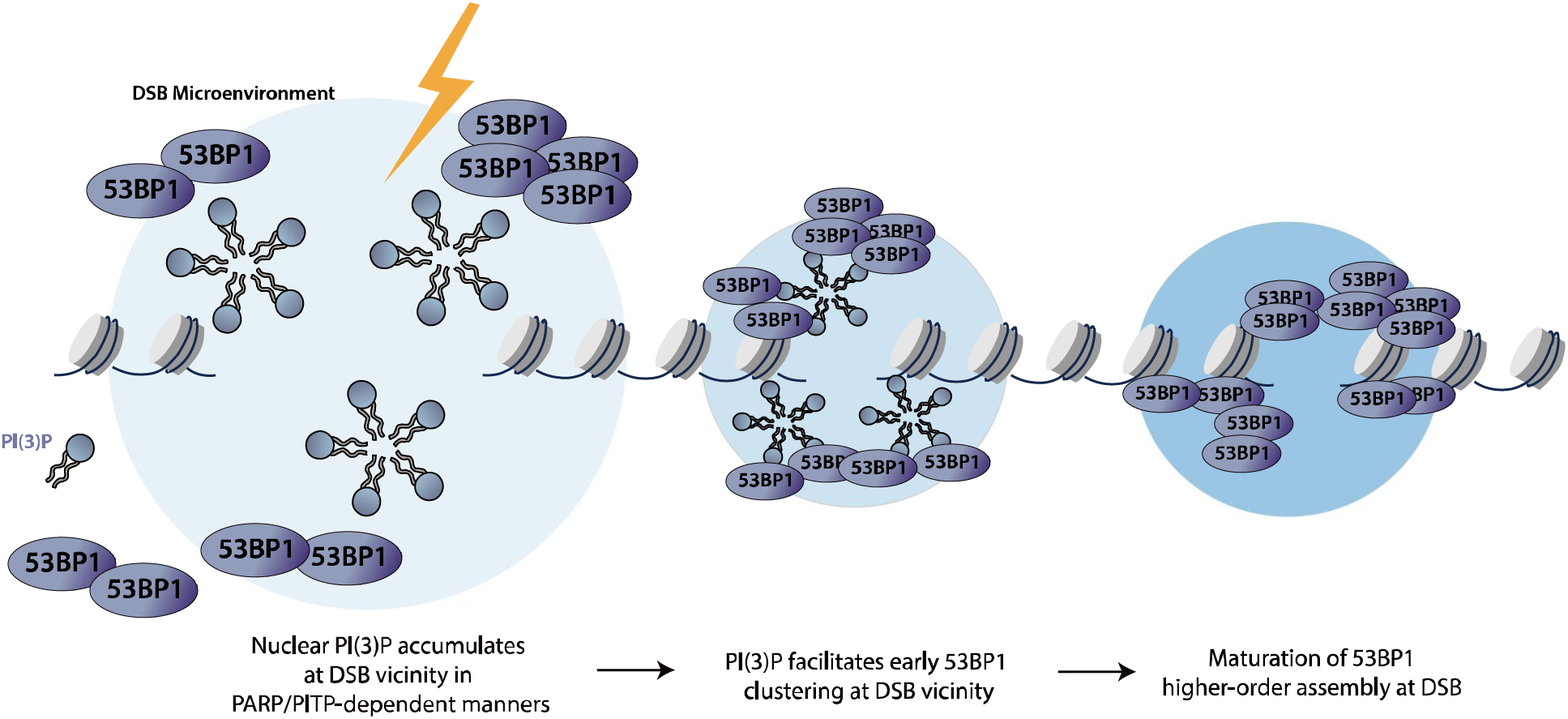
Working Model. Schematic illustration depicting maturation of 53BP1 higher-order assembly following DNA damage. DNA damage triggers PARP/PITP-dependent local concentration of PI(3)P in the vicinity of DSBs. Local concentration of 53BP1 is promoted by tBRCT-dependent, multivalent interactions with PI(3)P, after which 53BP1 transitions to γH2AX-dependent stable assembly on damaged chromatin.

## Discussion

In this study, we describe an unanticipated function for the 53BP1 tBRCT domain as a phospholipid-binding module. We show that tBRCT interacts specifically with PI(3)P, but not with other phosphoinositide species, and that multivalency drives strong binding of 53BP1 to PI(3)P clusters. Perturbation of the tBRCT-PI(3)P interaction compromised 53BP1 condensation *in vitro* and optodroplet formation *in vivo*, underscoring a role for tBRCT in promoting higher-order assembly of 53BP1. Notably, PI(3)P-binding-inactivating mutations in the tBRCT domain inhibit 53BP1 condensation more strongly than complete tBRCT deletion. One possible explanation is that deletion of tBRCT compromises the structural integrity of 53BP1, thereby promoting non-specific associations that can mimic condensation and mask the loss of function. In contrast, single point mutations preserve structural integrity and thus more clearly reveal the specific role of PI(3)P in driving condensation.

Regardless of whether 53BP1 condensation operates through LLPS, one plausible function of tBRCT is to nucleate and stabilize condensation through specific recognition of PI(3)P clusters at DNA damage sites. Within full-length or foci-forming 53BP1 constructs, the tBRCT domain may function as a multivalent anchor that interconnects multiple interaction epitopes^10^. In addition to serving as a PI(3)P-binding module described here, this domain also directly interacts with several DDR-associated factors, including γH2AX in a phospho-dependent manner^11^ and p53 in a modification-independent manner^40, 41^. Through these interactions, the tBRCT domain of 53BP1 may bridge lipid-enriched DSB sites to the formation of chromatin-bound repair assemblies. In this view, PI(3)P would contribute to the DNA damage specificity of the condensation process and the recruitment of DDR factors.

In line with the intricate nuclear dynamics observed during the cellular response to genotoxic stress^42^, our finding that chemical inhibition of PITP1/2 attenuates accumulation of GFP-p40PX at laser-induced DNA damage tracks suggests that DNA damage is coupled to PITP1/2-dependent transport of PI(3)P to the DSB microenvironment. PITP1/2 may help maintain the nuclear pool of this phosphoinositide, enabling its interaction with 53BP1 and thereby promoting higher-order assembly of 53BP1. In agreement with this model, sequestration of PI(3)P suppresses the formation of 53BP1 nuclear bodies prior to their stable accumulation on the damaged chromatin, highlighting an early role of PI(3)P in mobilizing 53BP1.

Although speculative, the recently described “compact” and “amorphous” classes of 53BP1^28^ may represent assemblies at distinct stages with different biophysical properties, including variations in mobility and foci morphology. Our observation that early 53BP1-containing DNA damage foci are particularly sensitive to 1,6-HD treatment suggests that these structures may correspond to initial nucleation events in the vicinity of DSBs, preceding stable assembly at γH2AX-decorated chromatin. While small non-coding RNAs generated at DSBs have been proposed as potential nucleators of 53BP1 condensation, our identification of PI(3)P as a specific and effective **promoter** of 53BP1 condensation highlights the multifaceted control of this key DDR factor. In summary, our work reveals an unexpected role for the 53BP1 tBRCT domain as a phospholipid-binding module and implicates phospholipid homeostasis as an important driver of 53BP1-dependent DDRs.

## Methods

### Cell culture and reagents

U-2 OS (ATCC, HTB-96) and their derivatives, and HEK 293T (ATCC, CRL-11268) cell lines were cultured in Dulbecco’s Modified Eagle Medium (DMEM) supplemented with 10% fetal bovine serum (FBS) and 1% penicillin/streptomycin. hTERT RPE-1 53BP1 knock-in cells^28^ were cultured under identical conditions at 37 °C with 5% CO_2_ supplied in the cell incubator. All reagents and resources used are listed in Supplementary Table 1.

### CRISPR Cas9 mediated gene editing

Guide RNAs (sgRNAs) targeting gene(s) of interest were designed, annealed, and cloned into the lentiviral vector lentiCRISPRv2 (Addgene #52961, Addgene #52962) according to the Zhang lab protocol. For lentivirus production, the lentiCRISPRv2-sgRNA plasmid was co-transfected with the packaging plasmids psPAX2 (Addgene #12260) and pMD2.G (Addgene #12259) into HEK 293T cells at a mass ratio of 4:3:1, using polyethylenimine (PEI) as the transfection reagent. Viral supernatant was harvested 48 hours post-transfection, passed through a 0.45 µm syringe filter (Sartorius), and immediately used for transduction or stored at -80 °C. Target cells were transduced with filtered viral supernatant in the presence of 8 µg/mL polybrene. At least 24 hours post-transduction, transduced cells were selected with 2 µg/mL puromycin (or blasticidin, as appropriate) for a minimum of 96 hours. The knockout efficiency was validated by Western blotting.

### Western blotting

Cells were collected and lysed with NETN buffer (150 mM NaCl, 1 mM EDTA, 20 mM Tris-HCl pH 8.0, 0.5% NP-40) with the addition of 25U/mL Benzonase nuclease and 1 mM MgCl_2_ for 20 minutes on ice. Cell lysates were then boiled with SDS loading buffer at 88 °C for 5 minutes followed by loading to SDS-polyacrylamide gel electrophoresis (SDS-PAGE) gel. Samples were then transferred to polyvinylidene fluoride (PVDF) membranes, and blocked by 3% skimmed milk in Tris-buffered saline with 0.1% Tween 20 detergent for at least 30 minutes. Membranes were then washed with TBST buffer 3 times for 5 minutes and then incubated overnight at 4 °C with indicated primary antibodies diluted in blocking buffer, after which membranes were washed again as above and incubated for 1 hour at room temperature (RT) with appropriate horseradish peroxidase (HRP)-conjugated secondary antibodies. Proteins were detected using SuperSignal™ West Pico PLUS Chemiluminescent Substrate, and chemiluminescent signals were captured using a ChemiDoc™ MP Imaging System.

### Immunofluorescence

Cells were seeded on 25 mm × 25 mm coverslips one day before fixation. Cells were washed once with phosphate-buffered saline (PBS). Unless otherwise mentioned, cells were fixed in 3% paraformaldehyde (PFA) for 20 minutes, and then permeabilized by 0.5% Triton X-100 for 10 seconds. Cells were then incubated with indicated primary antibody and fluorophore-conjugated secondary antibody diluted in 3% BSA. Cell nuclei were stained by DAPI diluted in PBS (1 µg/mL). Stained coverslips were mounted on glass slides with mounting medium. Images were then taken by Olympus BX53 microscope unless otherwise specified.

### Cell permeabilization

For experiments requiring RNase A or 1,6-hexanediol (1,6-HD) treatment, cells were first permeabilized. A solution of 1,6-HD (2% w/v) or RNase A (1 mg/mL) in 1% PBST was applied to live cells for 10 seconds at room temperature. Subsequently, cells were washed with PBS and immediately fixed with 3% PFA for 20 minutes, followed by standard immunofluorescence staining procedures as described above.

### Ionizing radiation

Cells were seeded on either 6-well plate (TPP Techno Plastic Products AG) or culture dish (TPP Techno Plastic Products AG) for different purposes on day before ionizing radiation (IR). Cells were subjected to IR treatment by cell irradiator at indicated dose, and following irradiation cells were processed for indirect immunofluorescence staining experimentations at time points specified.

### Laser micro-irradiation and live cell imaging by LSM 780

Cells were seeded on glass-bottom confocal dishes (SPL Life Sciences) and transfected with 4 µg of plasmid DNA. Approximately 24 hours post-transfection, laser micro-irradiation was performed using a Zeiss LSM 780 confocal microscope equipped with a 37 °C live-cell chamber maintaining 5% CO₂. A defined region within the nucleus of a transfected cell was subjected to DNA damage using a 750 nm two-photon laser, with power calibrated to deliver a localized dose of 12% of maximum output. Time-lapse imaging was initiated immediately using a 40x/1.4 NA oil DIC objective. Images were acquired every 2 seconds for the indicated duration using ZEN 2012 software. For quantification, the mean fluorescence intensity (MFI) within the micro-irradiated stripe was measured using ImageJ software. The relative recruitment of the fluorescent protein was calculated using the following formula:

MFI= MFI of micro-irradiation area - MFI of undamaged area

Relative MFI= (MFI of micro-irradiation area - MFI of undamaged area) / (MFI of micro undamaged area - MFI of background)

### 488 nm light induced phase separation in the Cry2 system

To induce optogenetic phase separation, cells were seeded on glass-bottom confocal dishes (SPL Life Sciences) and transfected with 4 µg of Cry2-fusion expression constructs.

Experiments were performed 24 hours post-transfection using a Zeiss LSM 780 confocal microscope equipped with a 37 °C live-cell chamber maintaining 5% CO₂. Phase separation was triggered by illuminating a defined cellular region with a 488 nm laser. For assembly kinetics, a continuous illumination at 0.5% of maximum laser power was applied for specified time duration. All images were acquired using a 40x/1.4 NA oil DIC objective and ZEN 2012 software (Carl Zeiss).

### Live super resolution microscope and 3D masking

Super resolution imaging was performed by live-SR microscope with a 100x oil immersion lens, and 3D masking is reconstructed by the Imaris software.

### Protein expression and purification

Constructs encoding maltose-binding protein (MBP)- and hexahistidine (6×His)-tagged protein of interest were transformed in BL21 bacteria two days before protein purification. A single transformed colony was inoculated into LB medium supplemented with the appropriate antibiotic and grown overnight at 37 °C. This starter culture was diluted into fresh medium and grown at 37 °C with shaking until the optical density at 600 nm (OD600) reached 0.6–0.8. Protein expression was induced with 1 mM isopropyl β-D-1-thiogalactoside (IPTG), followed by 16-hour overnight induction at 16 °C. Bacterial cells were collected and lysed in protein purification buffer (20 mM Tris-HCl (pH 8), 200 mM NaCl, 0.5% Triton X-100, 1 mM PMSF, 2 mM 2-Mercaptoethanol). Cell suspension was lysed by sonication on ice, and the lysate was centrifuged at high speed to yield clear supernatant. In parallel, amylose beads or Ni-NTA beads were washed with protein lysis buffer for at least 3 times.

Washed beads and the clear supernatant were co-incubated at 4 ℃ for at least 4 hours. Protein-bound beads were then washed with the lysis buffer again for at least 4 times. Bound proteins were eluted using elution buffer (20 mM Tris-HCl (pH 8), 200 mM NaCl, 1 mM PMSF, 2 mM 2-Mercaptoethanol) supplemented with either 60 mM maltose (for MBP fusions) or 250 mM imidazole (for His-tagged fusions). Eluted protein concentrations were determined using the bicinchoninic acid (BCA) assay.

### *In vitro* droplet assay

GFP tagged proteins were purified sequentially via their N- and C-terminal MBP and 6×His tags respectively, as described in the previous section. Purified protein was diluted to a final concentration of 5 µM in phase separation buffer (10 mM HEPES pH 7.4, 150 mM NaCl, 0.1 mM EDTA, 2 mM DTT, 5% PEG-8000). Where indicated, BODIPY-TMR labeled phosphoinositides were included at 2.5 µM. The reaction mixture was transferred to a glass bottom 96-well plate (Thermo Scientific), incubated for 20 minutes at room temperature to allow droplet formation, and then immediately imaged. Images were acquired using a Zeiss LSM 780 confocal microscope equipped with a 40x/1.4 NA oil DIC objective, controlled by ZEN 2012 software.

### Generation of giant unilamellar vesicles (GUVs)

A lipid mixture containing 20% of the specified phosphoinositide (e.g., PI(3)P) and 80% 1,2-dioleoyl-sn-glycero-3-phosphocholine (DOPC) was prepared in chloroform in a glass vial.

The organic solvent was evaporated under a gentle stream of nitrogen gas while rotating the vial to form a uniform lipid film on the inner surface. Residual solvent was removed by placing the vial under vacuum for at least 30 minutes. The dried lipid film was hydrated with 0.5 M sucrose solution (pre-warmed to 50 °C) to a final lipid concentration of 0.5 mM.

### MicroScale Thermophoresis (MST)

To assess the interaction between a recombinant protein (e.g., MBP-53BP1-C-6×His) and BODIPY-TMR-labeled phosphoinositide (e.g., PI(3)P), microscale thermophoresis (MST) assays were performed using a Monolith X instrument (NanoTemper Technologies). A constant concentration of 100 nM BODIPY-TMR phosphoinositide was incubated with serially diluted recombinant protein (starting at 15 µM, 1:1 dilution) in MST standard buffer (50 mM Tris-HCl, pH 7.4, 150 mM NaCl, 10 mM MgCl₂) to generate 16 concentration points. After a 15-min incubation at room temperature, samples were loaded into standard capillaries (Product No. MO-K022; NanoTemper Technologies) and analyzed at 25 °C with 100% excitation power and medium MST power (40%). Binding curves were fitted using the *K*_d_ model in Affinity Analysis v2.3 software (NanoTemper Technologies GmbH, München, Germany), and graphs were generated with GraphPad Prism 6. Fnorm(‰) was calculated by dividing F1 by F0. F1 is the fluorescence measured in the heated state, whereas F0 is the fluorescence measured before laser activation. The normalized Fnorm is plotted against the lowest ligand concentration Fnorm value, so that the coordinate curves start from 1.

### 53BP1 foci characterization and quantification

Foci were quantified using ImageJ (version 2.16.0). Images were background-subtracted, Gaussian-blurred, and segmented using auto local threshold with watershed separation. Foci measurements were extracted using Analyze Particles. Data from seven fields of view per condition were combined using Python (pandas) and visualized in R (ggplot2). Live-cell sequences (10-min intervals, 4–5 h) were tracked using TrackMate plugin. Fusion and fission events required validation over ≥3 consecutive frames (≥30 min). Event frequencies were normalized per cell and cross-validated with NIS Elements. Data were plotted in GraphPad Prism 10. Colocalization was analyzed using a custom ImageJ macro. Nuclei were segmented from 53BP1 channel after image pretreatment. Nuclei were detected using Measure Particles. 53BP1 foci were identified using adaptive thresholding. Colocalization was determined by measuring γH2AX intensity at each 53BP1 focus position, with positive scoring when intensity exceeded threshold (>15 arbitrary units). Three independent replicates were analyzed.

### Computational modeling

Phosphatidylinositol (PI) and seven phosphatidylinositide variants, differing in the number and positions of phosphate groups on the inositol ring, were generated in Avogadro^43^ and docked to the 53BP1 tandem BRCT (tBRCT) domain using AutoDock Vina v1.2.3^35, 36^, with a docking box size of 15 × 20 × 15 Å centered at the phosphate binding pocket. The 53BP1 tBRCT crystal structure used for docking corresponds to chain D of Protein Data Bank entry 5ECG^11^. Because phosphatidylinositides contain two long aliphatic chains that greatly increase conformational freedom and complicate docking, short-chain analogues were used (**Extended Data Fig. 4a**). This simplification is justified because binding is primarily driven by headgroup interactions.

For each ligand, the highest-ranked Vina docking pose was subjected to molecular dynamics (MD) simulations using the AMBER24 software suite^37^. For each system, five independent 1 μs replicate trajectories were generated. All systems were solvated in a cubic water box with at least 10 Å padding, and neutralized by adding sodium counterions. Each complex was first energy-minimized for 5000 steps using the steepest descent method, followed by gradual heating to 300 K in the NVT ensemble with a Langevin thermostat (γ = 2 ps⁻¹) over 100,000 steps. The systems were then equilibrated to 1 atm in the NPT ensemble using a Berendsen barostat^44^ over 500,000 steps. Heavy atoms were restrained with a force constant of 2.0 kcal·mol⁻¹·Å⁻² only during the heating and pressure-equilibration stages. An additional 10 ns of unrestrained equilibration was performed before the production runs. A 2 fs time step was used throughout, and bonds involving hydrogen were constrained using SHAKE ^45^. The ff14SB force field^46^ was used for the protein, and GAFF2^47^ was used for the ligands, with AM1-BCC charges assigned using ANTECHAMBER^47^. The TIP3P water model was used for all simulations^48^. Long-range electrostatics were treated using the particle mesh Ewald (PME) method^49, 50^, and van der Waals interactions were truncated at a 10 Å cutoff.

Structures extracted from the final restart files of the most stable 1 μs complex trajectories, together with structures from 100 ns MD simulations of the free ligands, were used as starting conformations for absolute binding free energy calculations using a stepwise thermodynamic integration (TI) approach^51^. Because phosphatidylinositide ligands are highly negatively charged, a two-step TI scheme was employed in which electrostatic and van der Waals interactions were decoupled separately. In addition, a softcore potential^52^ with parameters α = 0.2 and β = 50 Å^2^ ^53^was applied where α controls the degree of smoothing of the van der Waals potential and β modulates short-range distance scaling, thereby preventing singularities and large energy fluctuations as interactions are gradually turned off.

Except for TI-specific settings, the simulation parameters were the same as those used in the MD protocol. Twelve λ windows derived from Gaussian quadrature (0.00922, 0.04794, 0.11505, 0.20634, 0.31608, 0.43738, 0.56262, 0.68392, 0.79366, 0.88495, 0.95206, 0.99078) were used for alchemical electrostatic decoupling and van der Waals annihilation. The corresponding weights (0.02359, 0.05347, 0.08004, 0.10158, 0.11675, 0.12457, 0.12457, 0.11675, 0.10158, 0.08004, 0.05347, 0.02359) were used for numerical integration of ⟨∂U/∂λ⟩ to obtain the free energy change (ΔG). A 10 Å nonbonded cutoff was used, and a softcore potential^52^ was employed with α = 0.2 and β = 50 Å^2^ ^53^. Boresch-type six-degree-of-freedom (6-DoF) restraints were applied during the TI calculations. Each λ window was equilibrated for 200 ps in NVT, followed by 200 ps in NPT, and then a 10 ns production run was performed, with the final 6 ns used for TI analysis.

### Computational analysis of 53BP1 protein properties

The intrinsically disordered region and lipid binding prediction was done by the following websites respectively: https://iupred2a.elte.hu/^32^ and http://biomine.cs.vcu.edu/servers/DisoLipPred/^33^. Graphs were generated with GraphPad Prism 6.

### Statistical analysis and reproducibility

Unless otherwise specified, data from at least three independent biological replicates are presented as the mean ± standard error of the mean (SEM). Statistical analyses and graph generation were performed using GraphPad Prism software (version 6). For comparisons between two experimental groups, a two-tailed, unpaired Student’s t-test was applied. For comparisons involving more than two groups, one-way ANOVA with an appropriate post-hoc test was used. Differences were considered statistically significant at P < 0.05. Significance is denoted as follows: ns (not significant), *P < 0.05, **P < 0.01, ***P < 0.001, and ****P < 0.0001.

## Acknowledgements

We are grateful to Matthias Altmeyer for CRY2-mCherry-53BP1 expression constructs. This work is supported in part by funds from Research Grants Council (RGC-GRF No. 17103522 & 17101223) and Jessie Ho Professorship to MH.

## Author contributions

NX and MH conceptualized the study. NX performed all experiments unless stated otherwise with help from XX and YX. JT, SL and VD provided critical reagents. LY and CY shared expertise and provided supervision for biophysical and GUV assays. GC and GM performed the computational analyses. NX, MH, GC, and GM drafted the manuscript with input from all authors.

## Extended Data Figure Legend

**Extended Data Fig. 1 | Sorbitol treatment inhibits 53BP1 condensation.**

**a**, U2OS cells were fixed and stained with indicated antibodies post IR. Percentage of cells with non-γH2AX-overlapping 53BP1 nuclear bodies was quantified and plotted across three independent experiments (n = 300). Data are shown as mean ± SEM. Statistical significance indicated as ***P<0.001, ****P<0.0001; ns, not significant.

**b**, Sorbitol treatment preferentially suppresses early γH2AX-independent 53BP1 clustering events in RPE-1 GFP-53BP1 WT KI cells. Fluorescence intensity along the white line was quantified, and nuclei are outlined with dotted lines. Number of cells with more than 10 foci was also quantified and plotted.

**Extended Data Fig. 2 | Treatment with 1,6-hexanediol impairs H2AX-independent 53BP1 nuclear bodies.**

**a**, In H2AX-deficient U2OS cells, 53BP1 foci were visualized and quantified at 15 min and 120 min post 3 Gy IR with or without treatment with 1,6-HD (2% in 1% PBST). Nuclei are outlined with dotted lines. Quantification of cells with more than ten 53BP1 foci across three independent experiments was plotted (n = 300). Data are shown as mean ± SEM. Statistical significance indicated as ***P<0.001, ****P<0.0001; ns, not significant.

**b**, Live tracking of GFP-53BP1 fluorescence in RPE-1 cells by Ti-2E microscopy showed that 53BP1 foci area increased sharply during early cellular responses to bleomycin treatment. The line plot depicts the temporal changes in GFP-53BP1 foci number (blue) and area (grey).

**Extended Data Fig. 3 PI(3)P and RNA additively promote 53BP1 LLPS *in vitro*.**

**a-c**, PI(3)P and RNA ligand enhanced phase separation of MBP-GFP-53BP1-C-6×His proteins. *In vitro* phase separation assay was conducted with indicated purified proteins (5 µM) co-incubated with 2.5 µM BODIPY-TMR labeled PI(3)P and/or 2.5 µM RNA ligand. Results from quantifications from three independent experiments (n = 3) were plotted (c). Data are presented as mean ± SEM, with ****P<0.0001 indicating significant differences; ns, no significant difference.

**Extended Data Fig. 4 | Docking and molecular dynamics simulations of phosphatidylinositol and phosphoinositides interaction with 53BP1 tBRCT domain.**

**a,** Short-chain variants of phosphatidylinositol used for docking and free energy calculations. **b,** Top-scoring docking poses of phosphatidylinositol and phosphoinositides mapped onto the 53BP1 tBRCT domain. Phosphoinositides adopt a similar binding mode, with one phosphate group occupying the canonical phosphate-binding pocket, including residues K1773 and K1814.

**c,** Stability of five independent 1 μs MD simulations for phosphatidylinositol and phosphoinositides. RMSD values were calculated by fitting the heavy atoms of the 53BP1 tBRCT domain. Each trace represents an independent replica.

## Supplementary information

**Supplementary Table 1.**
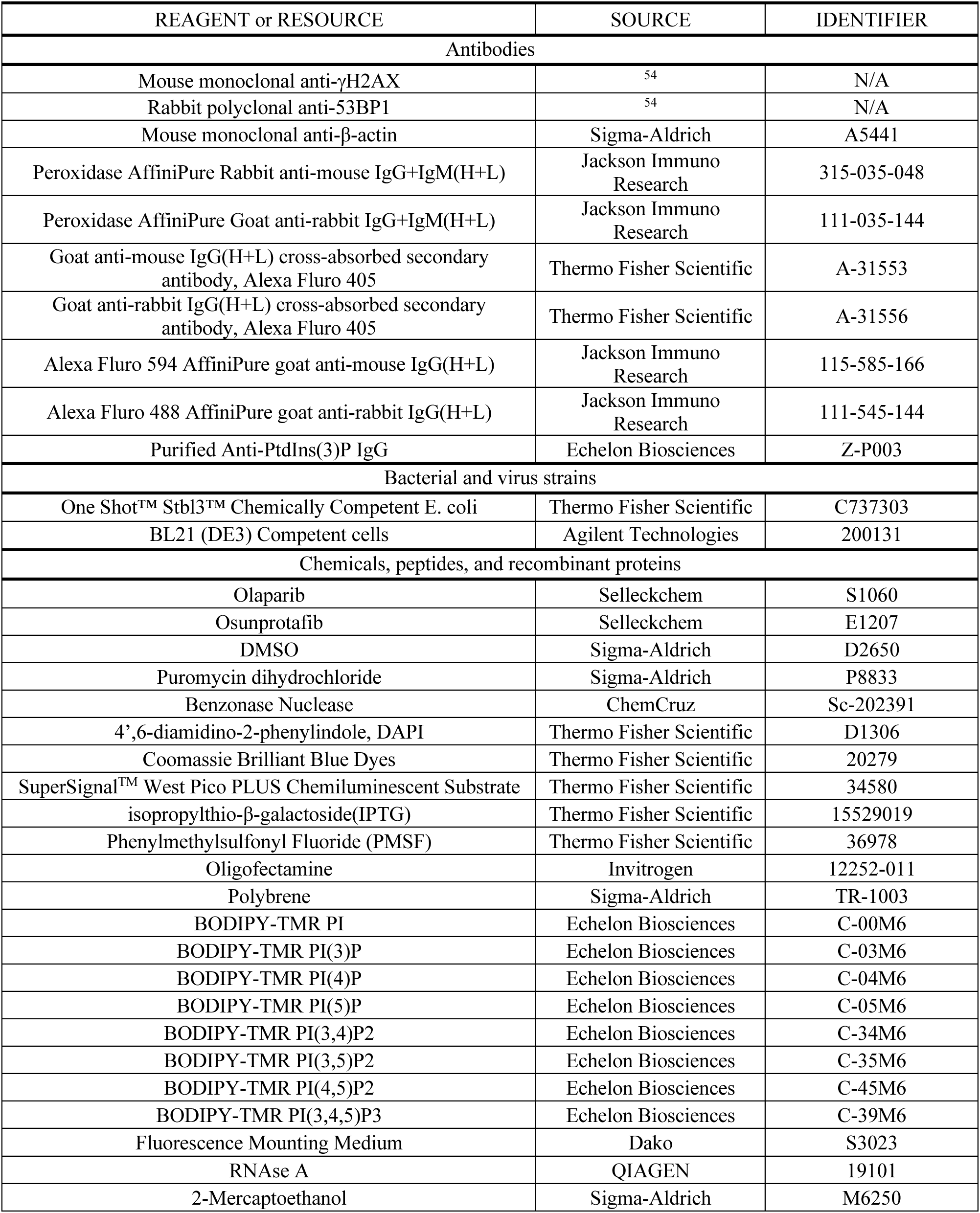
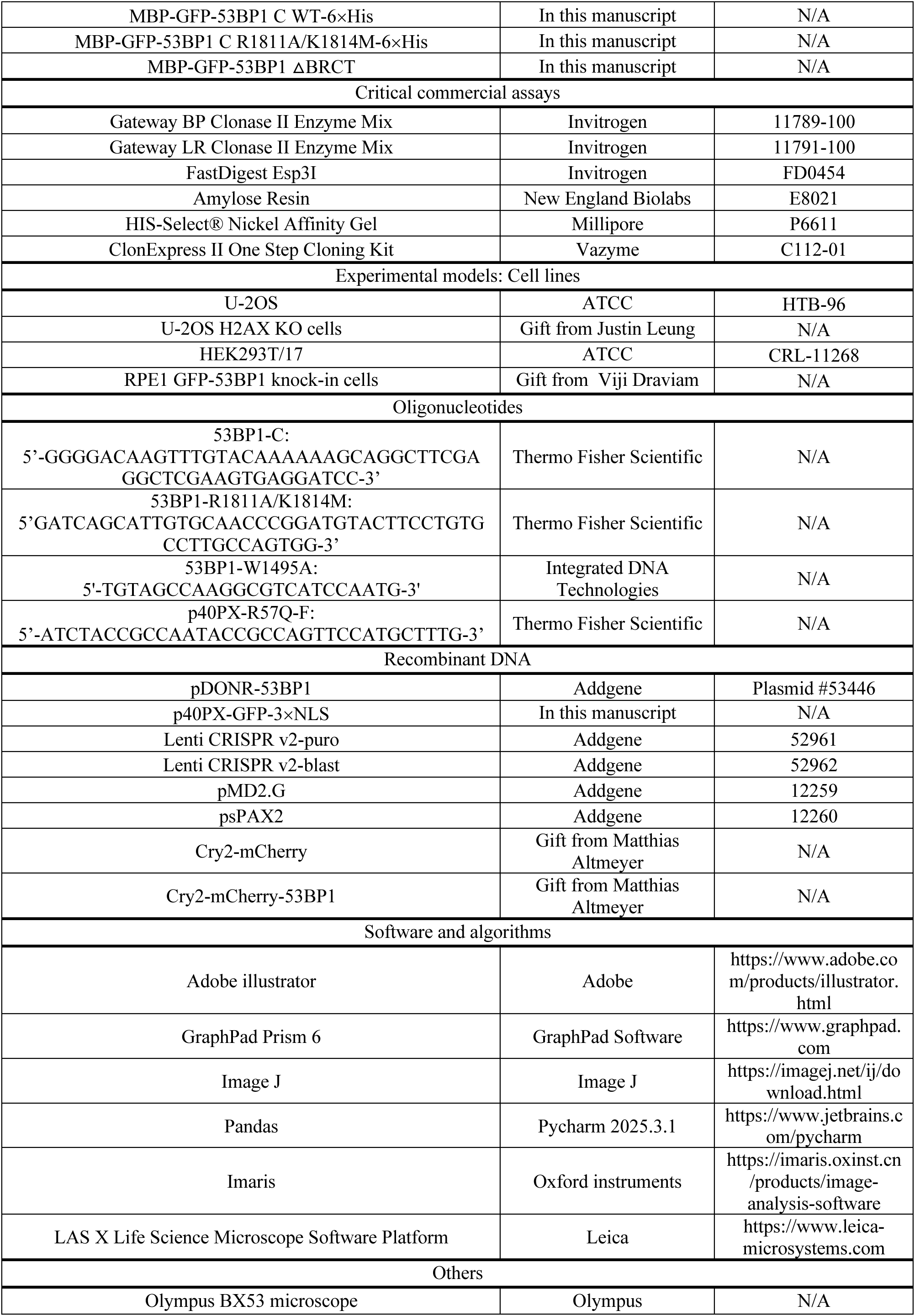
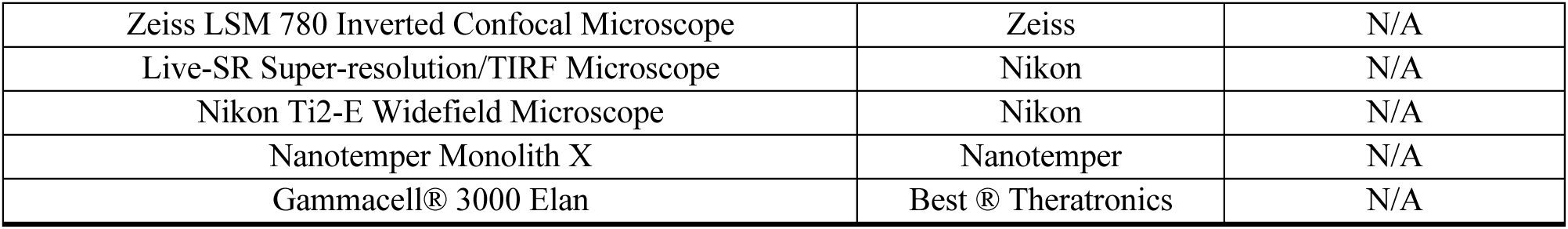
List of Reagents and Resources.

