## Supplementary figures and images for "Phosphatidylinositol 3-phosphate promotes 53BP1 condensate-like assembly at DNA double-strand breaks"

### extended data figures

#### Extended Data Figure 1

a

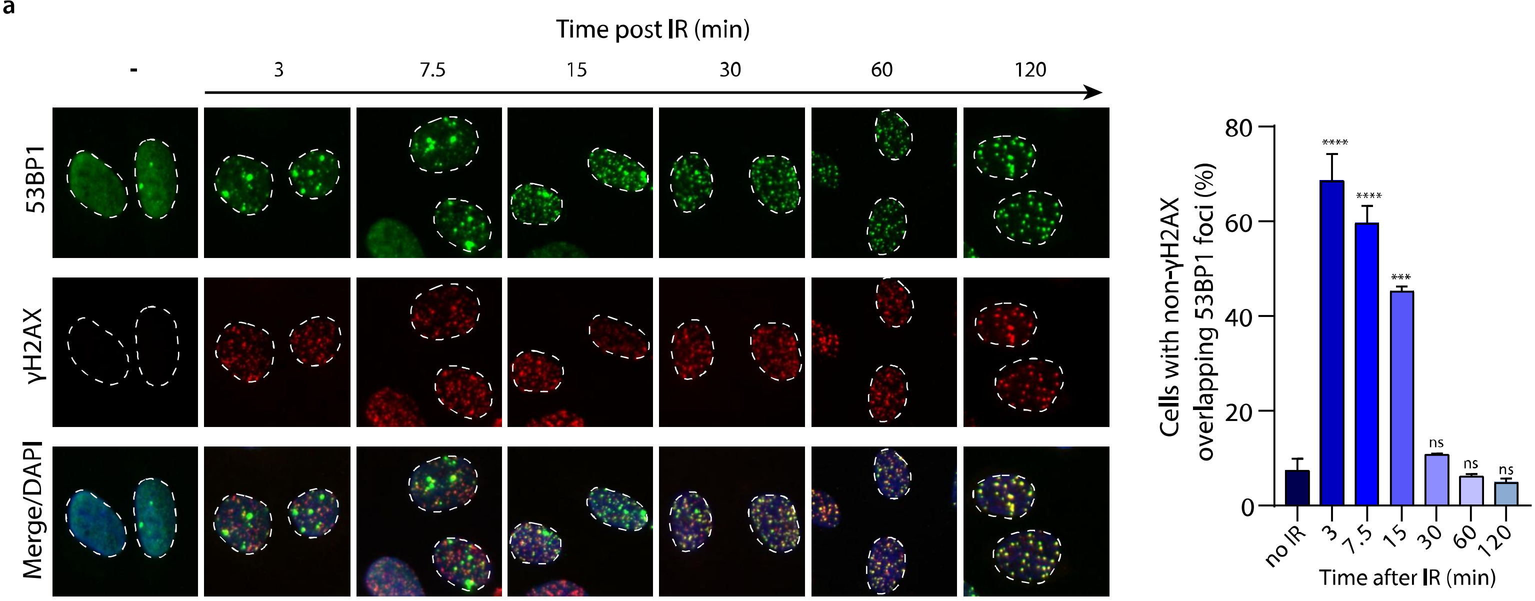


b

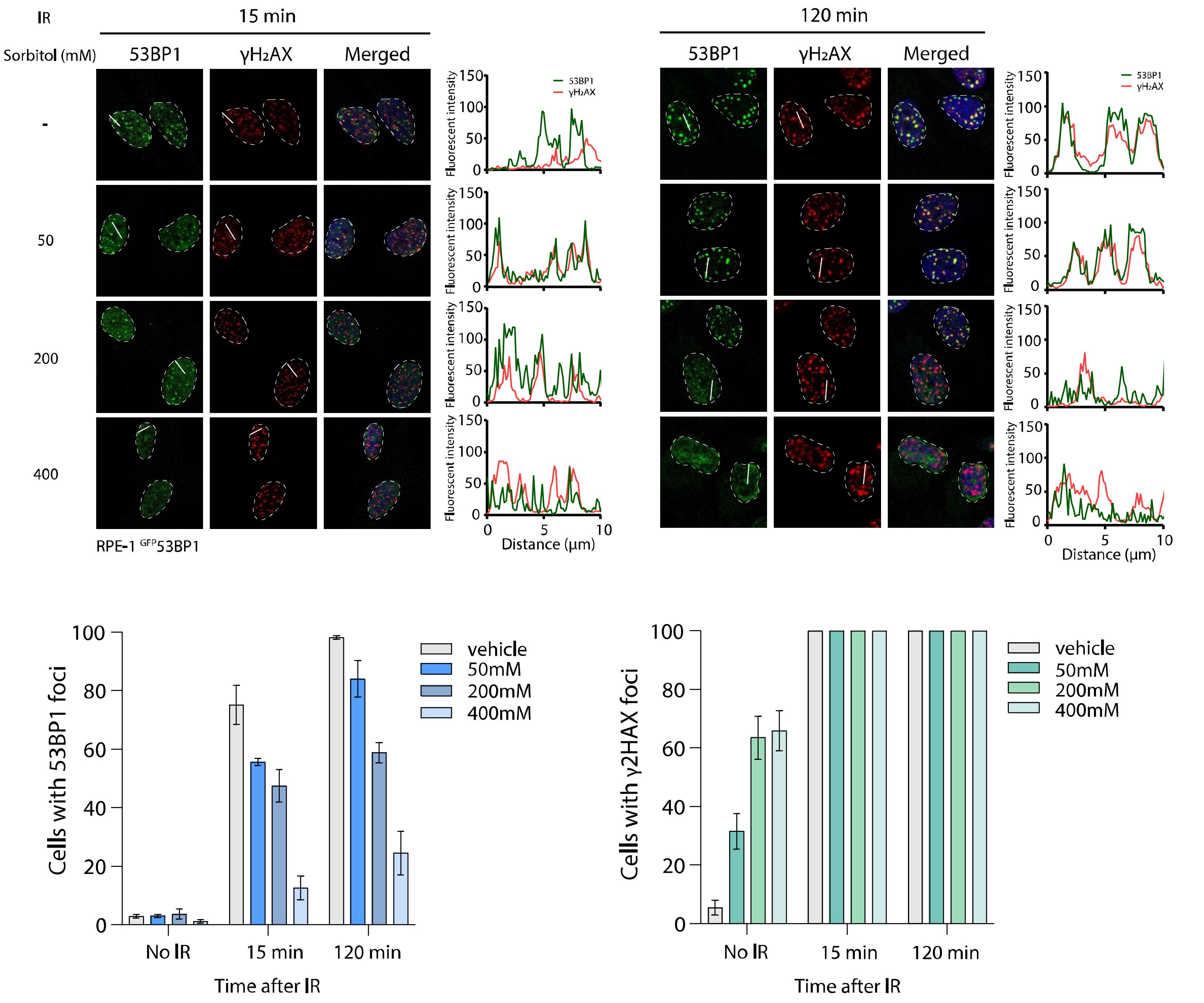

**Extended Data Figure 2**

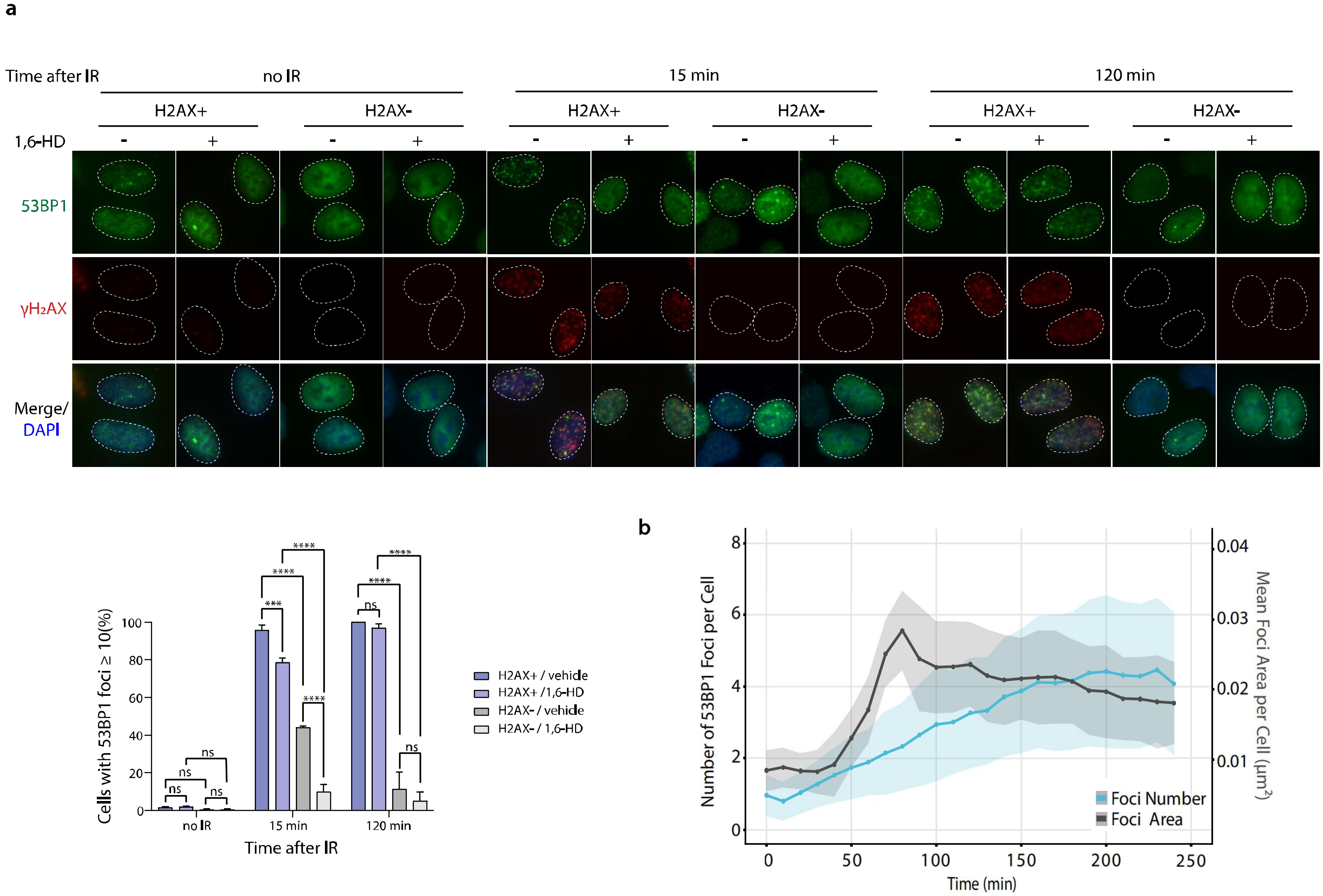

##### Extended Data Figure 3

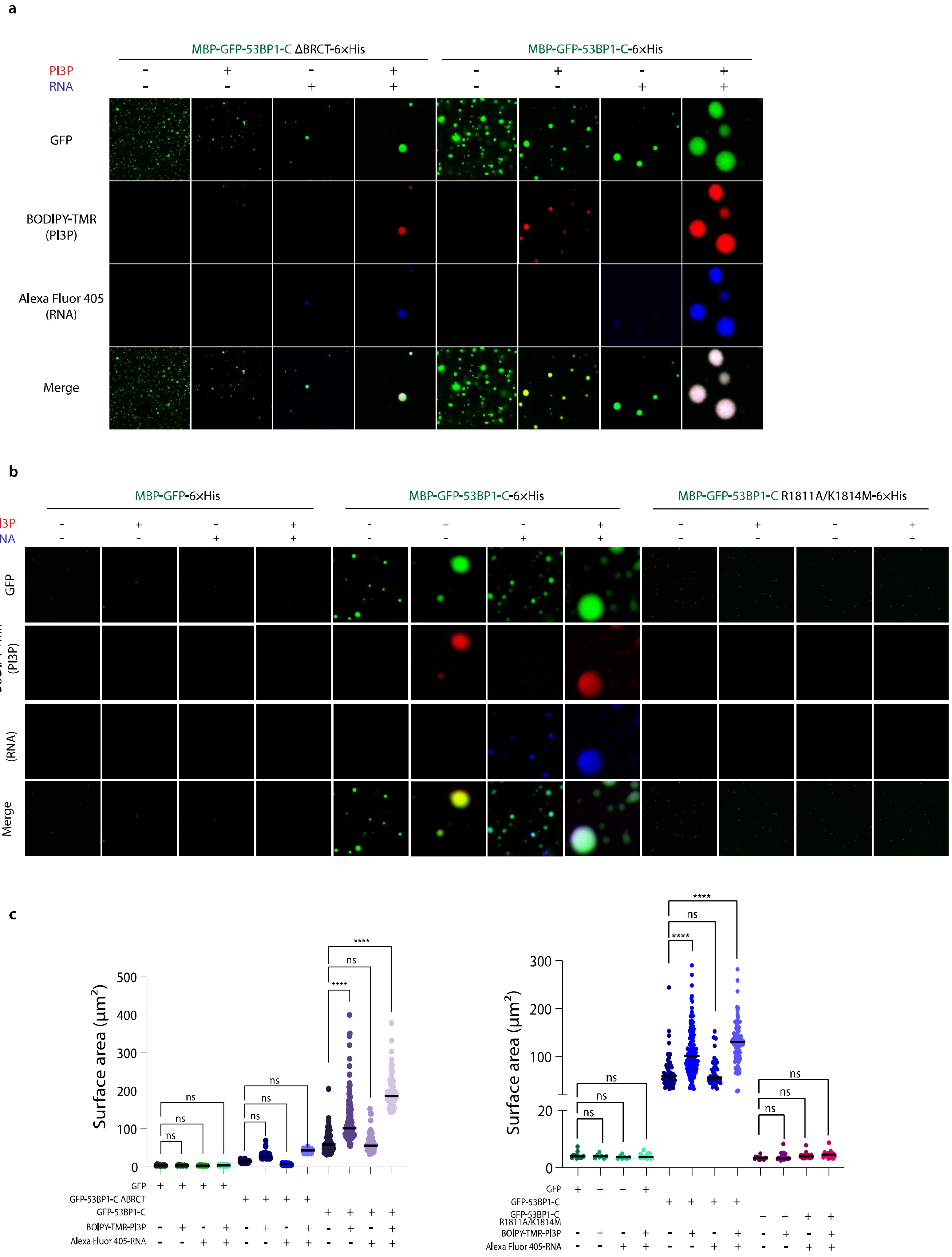

#### Extended Data Figure 4

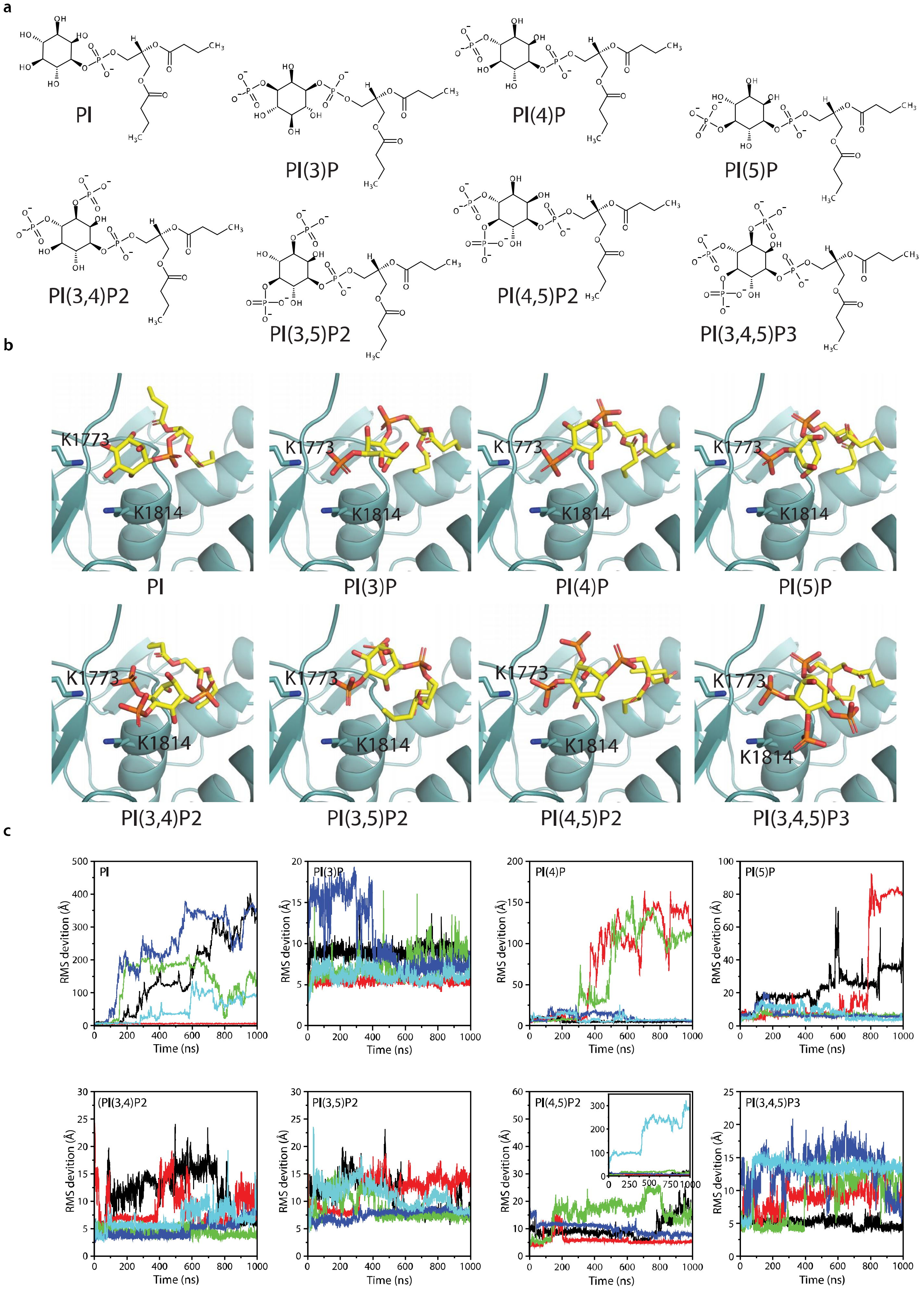
